# Dispensing a synthetic green leaf volatile to two plant species in a common garden differentially alters physiological responses and herbivory

**DOI:** 10.1101/370692

**Authors:** Grace E. Freundlich, Maria Shields, Christopher J. Frost

## Abstract

Herbivore-induced plant volatile (HIPV)-mediated eavesdropping by plants is a well-documented, inducible phenomenon that has practical agronomic applications for enhancing plant defense and pest management. However, as with any inducible phenomenon, responding to volatile cues may incur physiological and ecological costs that limit plant productivity. In a common garden experiment, we tested the hypothesis that exposure to a single HIPV would decrease herbivore damage at the cost of reduced plant growth and reproduction. Lima bean (*Phaseolus lunatus*) and pepper (*Capsicum annuum*) plants were exposed to a persistent, low-dose (∼10ng/hour) of the green leaf volatile *cis*-3-hexenyl acetate (*z3*HAC), which is an HIPV and damage-associated volatile. *z*3HAC-treated pepper plants were shorter, had less aboveground and belowground biomass, and produced fewer flowers and fruits relative to controls while *z*3HAC-treated lima bean plants were taller and produced more leaves and flowers than did controls. Natural herbivory was reduced in *z3*HAC-exposed lima bean plants, but not in pepper. Cyanogenic potential, a putative direct defense mechanism in lima bean, was lower in young *z3*HAC-exposed leaves, suggesting a growth-defense tradeoff from *z3*HAC exposure alone. Plant species-specific responses to an identical volatile cue have important implications for agronomic costs and benefits of volatile-mediated inter-plant communication under field conditions.

## Introduction

Production and utilization of airborne chemical cues is prevalent within the plant kingdom. Plants depend on airborne chemical signaling for pollination [1], indirect defense [2], protection from pathogens [3], and herbivore resistance [4]. Volatile communication is also pivotal for plant-plant signaling, and selection for such signaling depends on honest cues that reliably confer ecologically relevant information. For example, herbivory is a fundamental ecological interaction that impacts plant fitness, and many plants increase the production and emission of volatile compounds in response to herbivore damage [5]. Such herbivore-induced plant volatiles (HIPVs) are potentially reliable cues around which plant-plant eavesdropping could be evolutionarily adaptive [6]. Undamaged plants (or parts of the same plant [7,8]) eavesdropping on HIPVs from a plant experiencing herbivory may directly trigger stress responses [9-11], or alternatively prime responses for future potential herbivory [4,7].

HIPV-mediated eavesdropping appears to be a common phenomenon. For example, HIPVs prime or induce corn [12,13], tomato [14], poplar [4,7], blueberry [15] and lima bean [16,17] against herbivory. HIPVs can be diverse and taxa-specific [18,19], but are often comprised of monoterpenes, sesquiterpenes, benzenoids and green leaf volatiles GLVs [20,21]. In contrast to volatile terpenes and benzenoids [18,22], GLVs are immediately released into the airspace whenever leaves are mechanically damaged [23] serving as early indicators of wounding and herbivory. GLV exposure alters gene expression profiles related to specialized metabolite production and accumulated secondary metabolite precursors in preparation for inducing resistance [24]. For example, the GLV *cis*-3-hexenyl acetate (*z*3HAC) induces transcriptional changes in poplar [4] and maize [21] that prime oxylipin signaling and induced resistance. Like other GLVs, *z*3HAC can be emitted from wounded leaves alone, but may also represent a reliable cue because it is typically released from herbivore-damaged leaves in a variety of species [23], including tomato [14], maize [21], Arabidopsis [25], lima bean [8,17], pepper [26,27], and poplar [4,28].

Costs incurred by plants responding to airborne HIPV cues alone are largely unknown. Plant defense theory posits that induced resistance by plants against herbivores is a cost-savings strategy to restrict the deployment of costly specialized defensive metabolites until necessary [29,30]. However, inducible resistance generates a period of vulnerability between the time of attack and the upregulation of resistance [31]. Perception of early reliable cues may overcome such a vulnerability by allowing a plant to anticipate a probable attack and prime defenses before herbivory occurs. Since HIPV-mediated priming is an inducible phenomenon, theory predicts that responding to reliable cues alone should incur costs that outweigh and select against maintaining a “primed state” [32,33]. In other words, perception of a priming stimulus, such as an HIPV, may induce physiological changes that incur costs that are less expensive than induced resistance itself. Previous work with non-volatile priming agents β-amino butyric acid (BABA) [34] and snail mucus [35] both support this prediction. Similarly, costs associated with volatile perception alone that initiate priming should be less severe than costs of induced resistance to actual herbivory [36]. Yet there is currently limited experimental evidence of such costs with respect to anti-herbivore volatile cues. For example, bacterial-derived volatiles 3-pentanol and 2-butanone increased fresh fruit weight in field-grown *Cucumis sativa* [37], wild tobacco (*Nicotiana attenuata*) exposed to airspace of experimentally clipped sagebrush produce more seeds (i.e., higher presumptive fitness) relative to control plants [38]. These results suggest that ecological costs of exposure to volatile cues may be context dependent, but comparative cost/benefit tradeoffs for perception of HIPVs alone among sympatric field-grown plants i s currently lacking.

Here, we report a common garden field experiment with lima bean (*Phaseolus lunatus*) and chili pepper (*Capsicum annuum*) testing the hypothesis that field plants subject to a persistent dose of an ostensibly reliable volatile cue incur consistent costs reflected in reduced growth and reproduction. We treated individuals of both species to repeated low-dose applications of *z3*HAC and measured their growth, reproduction, and herbivore damage throughout the growing season. We predicted that exposure to *z*3HAC—regardless of plant species identity—would reduce growth and reproductive output, while also reducing natural herbivory.

## Materials and Methods

### Study Site and Plants

A common garden experiment was established on a 54m^2^ plot within Blackacre Conservancy’s community garden in Louisville, Kentucky (38°11’33.8“N 85°31’28.3”W; Supplemental Fig. 1). The field site was enclosed in a mesh fence to exclude mammalian herbivores. *Phaseolus lunatus*, Fabaceae, variety Fordham Hook 242 (‘lima bean’) and the *Capsicum annuum*, Solanaceae, variety Cayenne pepper, Joe Red Long (‘pepper’) were chosen as phylogenetically distinct model plants with previously established defense profiles [39,40]. Seeds were purchased from the Louisville Seed Company (Louisville, KY, USA), and germinated in Metromix 510© in May 2016 in the Biology Department’s greenhouse. After reaching ∼20cm in height, 132 lima bean plants were transplanted to the field May 30, 2016, at 4 weeks old and 98 pepper plants at 8 weeks old were transplanted to the field June 28, 2016. While both species were started at the same time, peppers were placed in the field later than the lime beans because they needed additional maturation time before transferring to the field. Within the field site, plants were planted in alternating rows of twos of lima bean and pepper. Distance limitations exist regarding plant volatile perception [41] and consequent herbivore resistance [7,42]. Previous studies with sagebrush [43] and lima bean [44] indicate that volatile cues are effective over relatively short distances of less than 100cm. Therefore, all plants in our experiment were spaced one meter apart from one another in all directions to reduce the risk of interplant communication and cue crossover.

### Volatile exposure manipulations

Plants were acclimated to the field for one week after planting before volatile treatments began. To simulate a naturally occurring low dose [45,46], plants were exposed to lanolin infused with the equivalent of 10ng/hr of *z*3HAC, a concentration 25% of that which previously primed poplar [4] and maize [21]. A treatment vial contained 50mg of a 30ng/ul *z*3HAC/lanolin, while a control vial contained 50mg of lanolin. Each glass vial had a 9mm aperture and was maintained at -80°C until use. Each week, both the *z*3HAC-infused lanolin vials and lanolin-only controls were placed at the bottom of their respective plants. Each vial was inverted and supported with a wire stand and each vial was wrapped in aluminum foil to reduce photodegradation [47] (Supplemental Fig. 2). These vials were left in the field and replaced every seven days for the duration of the field season. Plants were randomly assigned to either *z*3HAC treatment (lima bean n=63; pepper n=35) or lanolin control (lima bean n=72; pepper n=43). The unit of replication was an individual plant and each plant received its own vial. Random assignment of treatments was made using blocks of 4 adjacent plants; block was included as a random factor in statistical models, and was not a significant factor in any of the models.

### Growth, biomass, and reproduction measurements

We measured height and total leaf counts routinely on the experimental plants. Leaves were only counted if they were wider than 2cm across for both species while height measurements for both species were recorded from the base of the plant to the uppermost branching point. For lima bean, height was determined by measuring the longest runner within the bush, while pepper plants were measured from the base of the main stalk to the highest branching point. Along with height, the total number of leaves per plant was measured throughout the field season. A complete biomass harvest was conducted on pepper for leaves, roots, and stems at the end of the field season. All leaves and fruits were separated into paper bags before individual plants were extracted from of the ground. After removal, roots and stems were separated, roots were washed with water to remove dirt, and placed into separate paper bags. All materials were dried at 60°C for 24 hours and then weighed. A biomass harvest for lima bean was not performed because an *Epliachna varivestis* (Mexican Bean Beetle) outbreak late in the season removed much of the leaf tissue before we could determine reliable biomass measurements.

We measured total flower and fruit production in both species. Flowers were recorded if they were true flowers with fully mature pistils and stamen. If a flower was not fully mature, it was recorded as a flower bud. Fruits were recorded as soon as fruit development was observed with either initial pod or exocarp development. Throughout the field season, fruit and flower counts per plant were recorded along with the number of mature and immature fruits.

From the fruits harvested from the final biomass harvest, ∼10 randomly selected, mature fruits from each pepper plant were chosen for seed count analysis (188 fruits from z3HAC-treated plants and 210 fruits from controls). Dried fruits were dissected with a scalpel and all seeds were isolated and counted.

### Herbivory

Leaf chewing damage was assessed for both pepper and lima bean as percent leaf area removed (LAR) using a visual estimation technique [48,49] with the following damage categories: 0%, 0-5%, 5-15%, 15-30%, 30-50%, 50-70%, 70-90%, and >90%. For each damage assessment, every leaf on a plant was categorized into one of the damage categories, and an overall percent damage was determined as a weighted average of all leaves. Plants were also routinely monitored for the presence of naturally occurring chewing and piercing/sucking herbivores. In particular, we observed an ephemeral, natural occurrence of the black bean aphid (*Aphis faba*), and recorded its presence/absence on lima bean plants in the field.

### Leaf collections and cyanide measurements

Since cyanogenic potential (CP) is an inducible herbivore defense in lima bean [50,51], we used CP as a metric for induced responses in the presence of *z3*HAC. We collected source and sink leaves [52,53] on July 10, 2016, which was approximately 6 weeks into the field season. We developed a novel protocol in the lab for microscale colorimetric CP quantification by modifying an existing macro-scale protocol [54,55]. Briefly, 5mg of lyophilized tissue was mixed with 200uL of citrate buffer (0.1 M, pH 5.5-6.5) in a 2mL centrifuge tube, into which a 200uL glass vial (Agilent #5183-2090) containing 100uL 1.0M NaOH was also placed and the centrifuge tube caped completely. After 15 hours, a 50uL aliquot of the 1.0M NaOH in the inner glass vial was diluted to 0.1M, and a 30uL aliquot was neutralized with 30uL 0.5M acetic acid in a 96-well reaction plate. Then, 75 μL of Reagent A (5 mg/ml succinimide [VWR, AAA13503] and 0.5 mg/ml n-chlorosuccinimide [VWR, AAA10310]) and then 30 μL of Reagent B (30 mg/ml barbituric acid [VWR, BT134930] in 30% pyridine [Sigma, 270970]) was added to each well. After 8 min, absorbance at 580nm was measured on a plate reader (Molecular Devices SpectraMax M2). Quantification of CP was made against a standard curve of NaCN (VWR, BT212960).

### Statistical analyses

All statistical analyses were performed in R (version 3.4.2) with the lme4 and multcomp packages. Growth data, such as plant height, leaf area removed, and flower counts, were analyzed using repeated measures ANOVA with the aov function with a Gaussian distribution. For repeated measures analyses, we treated date as a within-subjects effect and treatment as a between-subjects effect for all analyses. Differences between treatments at each individual time point, as well as all biomass data, were analyzed using one-way ANOVA (glmer function) followed by a Tukey’s post hoc comparison. Cyanogenesis data were log transformed to satisfy assumptions of normality before being analyzed by a glm model followed by a Tukey’s post hoc comparison.

## Results

### *z*3HAC differentially effects growth of lima bean and pepper plants

Treatment with *z3*HAC differentially affected the growth of lima bean and pepper plants. On average, *z*3HAC-treated lima bean grew 11% taller compared to control plants throughout the field season (Fig.1A; F_2,927_=9.688, *P*=0.002) and produced 17% more leaves overall than did controls (Fig.1B; F_1,571_=4.339, *P=*0.038). In contrast, *z*3HAC-treated pepper plants were 12% shorter relative to controls (Fig.1C; F_1,48_=6.168, *P=*0.017) and produced 23% fewer leaves over the field season (Fig.1D; F_1,237_=21.58, *P<*0.001). Consistent with height and leaf counts, *z3*HAC treatment reduced overall biomass of pepper plants by 24% on average (Fig.2). When we destructively harvested all pepper plant biomass at the end of the season, *z*3HAC-treated pepper plants had lower leaf, stem, and root dry biomass by 21%, 31%, 29%, respectively (Fig.2A-C) (Z=3.379, *P<*0.001; Z=-2.035, *P=*0.042; Z=-2.379, *P=*0.017). Despite these *z*3HAC-mediated effects on biomass exposure, the aboveground-to-belowground biomass ratio was similar regardless of treatment (Fig.2D; Z=0.31, *P=*0.757). That is, pepper plants treated with *z*3HAC were smaller relative to control plants.

**Fig. 1.**
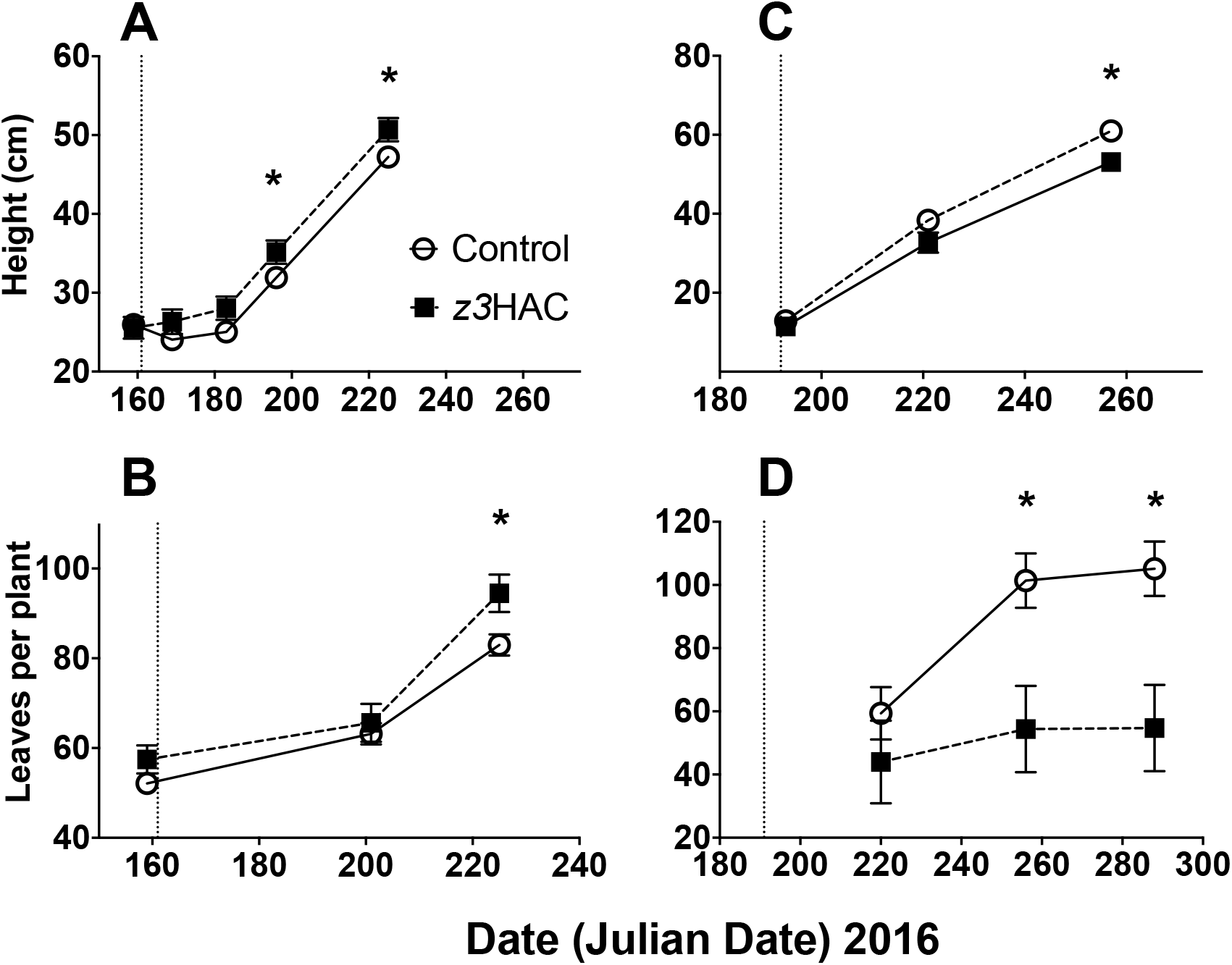
Height measurements and leaf counts for *Phaseolus lunatus* (lima bean) and *Capsicum annuum* (pepper) grown in a common garden field experiment. Height for (A) lima bean was measured from the base of the longest runner to the uppermost branching point; (C) pepper height was measured from the base of the main stalk to the uppermost branching point. Leaf counts for (B) lima bean and (D) pepper included all mature leaves on each plant. Open circles represent control plants (receiving lanolin-filled vials); filled squares represent plants receiving a persistent application of vials containing 10ng/hr *cis*-3-hexenyl acetate (*z*3HAC) dissolved in lanolin. Dropdown lines indicate the initial application of *z*3HAC treatment: lima bean and pepper plants were first exposed on June 10, 2016 (Julian date 161) and July 11, 2016 (Julian date 192), respectively. Points represent averages +/- SE. Repeated measures ANOVAs (aov in R) were followed by one-way ANOVAs at each time-point. Asterisks (*) represent *P*<0.05 between treatment and control at each time point. See Supplemental Table 1 for complete statistics.

### *z*3HAC reduces reproductive output in pepper plants

*z*3HAC treatment also differentially affected reproductive output between the two species, and lowered fruit output in pepper. Flower production was 30% higher in lima bean plants exposed to *z*3HAC (Fig.3A; F_1,576_=15.044, *P*<0.001), while *z*3HAC-treated peppers produced 37% fewer flowers relative to control plants at the end of the field season (Fig.3B; F_1,43_=14.48, *P*<0.001). *z*3HAC-treated pepper plants also produced 23% fewer fruits overall relative to controls (Fig.4A; Z=-2.035, *P=*0.042), and the fruits that were produced by *z*3HAC-treated plants had lower wet and dry masses (Fig.4B-C; Z=-2.88, *P=*0.004; Z=-2.439, *P=*0.015), and 10% lower total seed counts (Fig.4D; Z=3.524, *P<*0.001) and total seed masses (Fig.4E; Z=3.334, *P*<0.001), relative to controls. That said, the ratio of seed mass to fruit mass was similar between *z*3HAC-treated and control plants (Fig.4F; Z=0.588, *P=*0.807), as was mass of an individual pepper seed (Supplemental Fig.3). There was no apparent difference in lima bean pod production (Supplemental Fig.3), though an unexpected field-wide premature pod drop independent of treatment prevented us from fully determining lima bean pod/seed production with confidence.

**Fig. 2.**
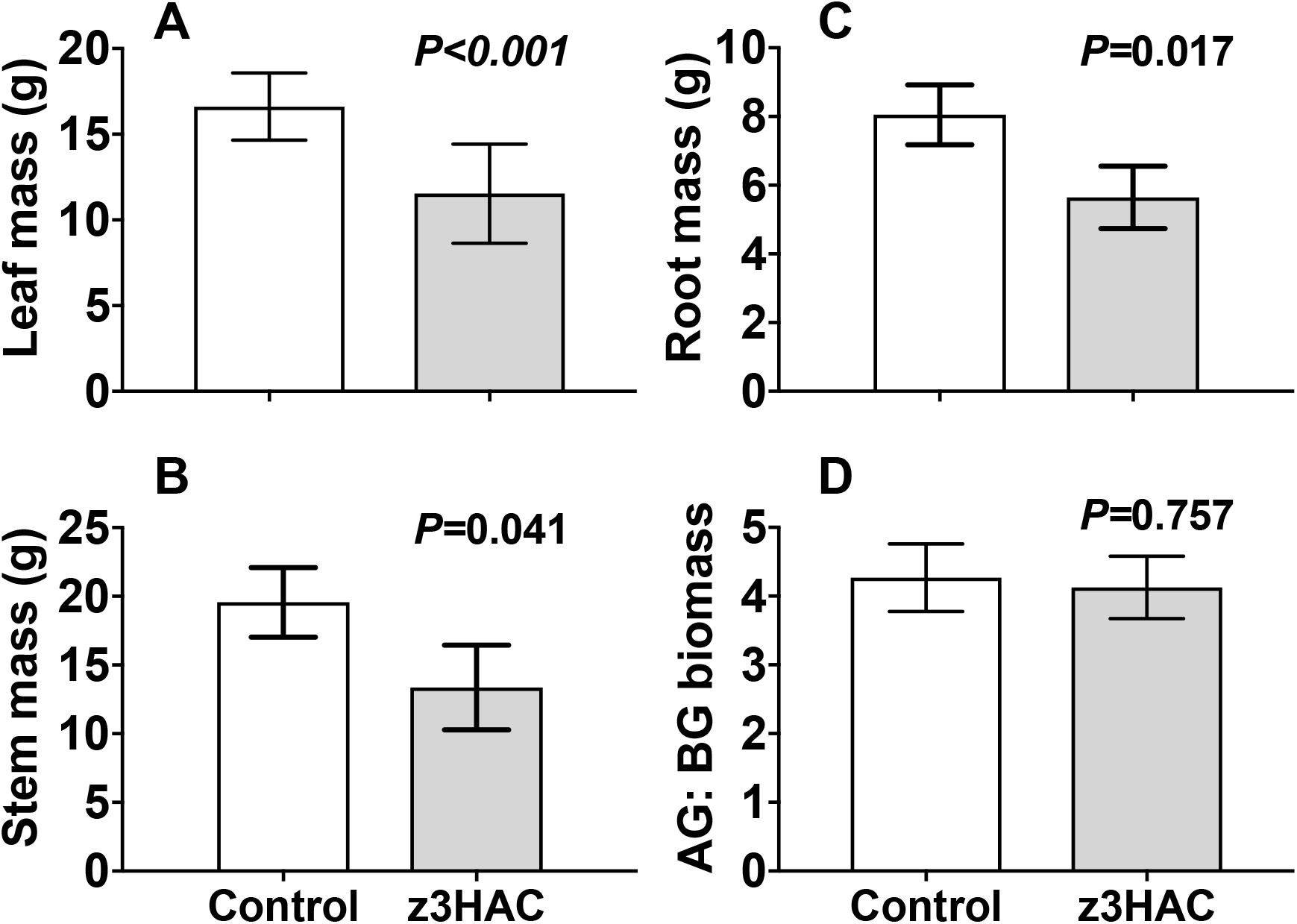
Biomass measurements of field-grown *Capsicum annuum* (pepper) plants. (A) Leaf, (B) stem, (C) root biomass, and (D) the aboveground:belowground biomass ratio in *C*.*annuum* plants were determined at the end of the field season following destructive harvest. Bars represent means +/- S.E.M.. *P*-values represent Tukey’s HSD comparisons. See Supplemental Table 2 for complete statistics.

**Fig. 3.**
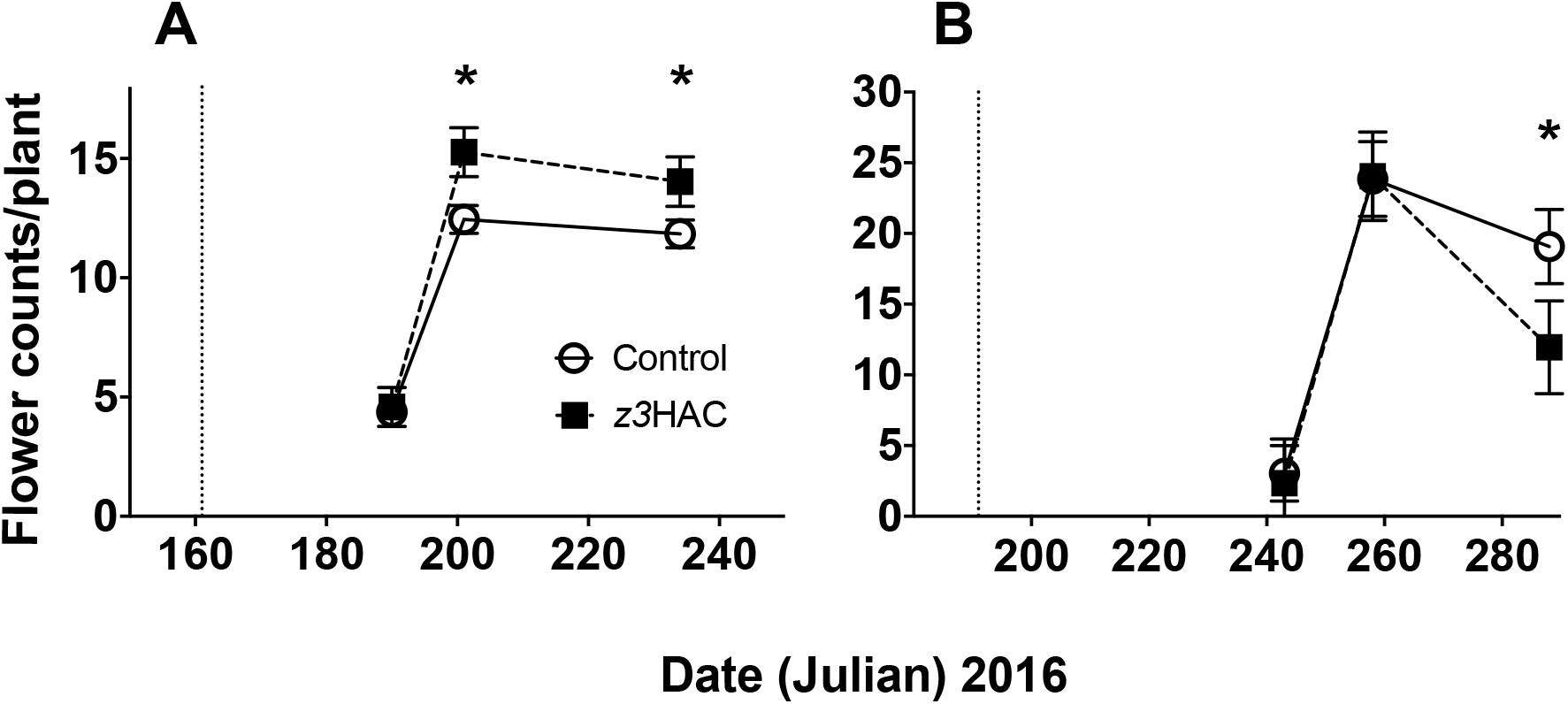
Total flower production in (A) *Phaseolus lunatus* (lima bean) and (B) *Capsicum annuum* (pepper). Open circles represent control plants (receiving lanolin-only vials); filled squares represent plants receiving a persistent application of vials containing 10ng/hr *cis*-3-hexenyl acetate (*z*3HAC) dissolved in lanolin. Dropdown lines indicate the initial application of *z*3HAC treatment: lima bean and pepper plants were first exposed on June 10, 2016 (Julian date 161) and July 11, 2016 (Julian date 192), respectively. Points represent averages +/- SE. Asterisks (*) represent *P*<0.05 between treatment and control at each time point. See Supplemental Table 3 for complete statistics.

**Fig. 4.**
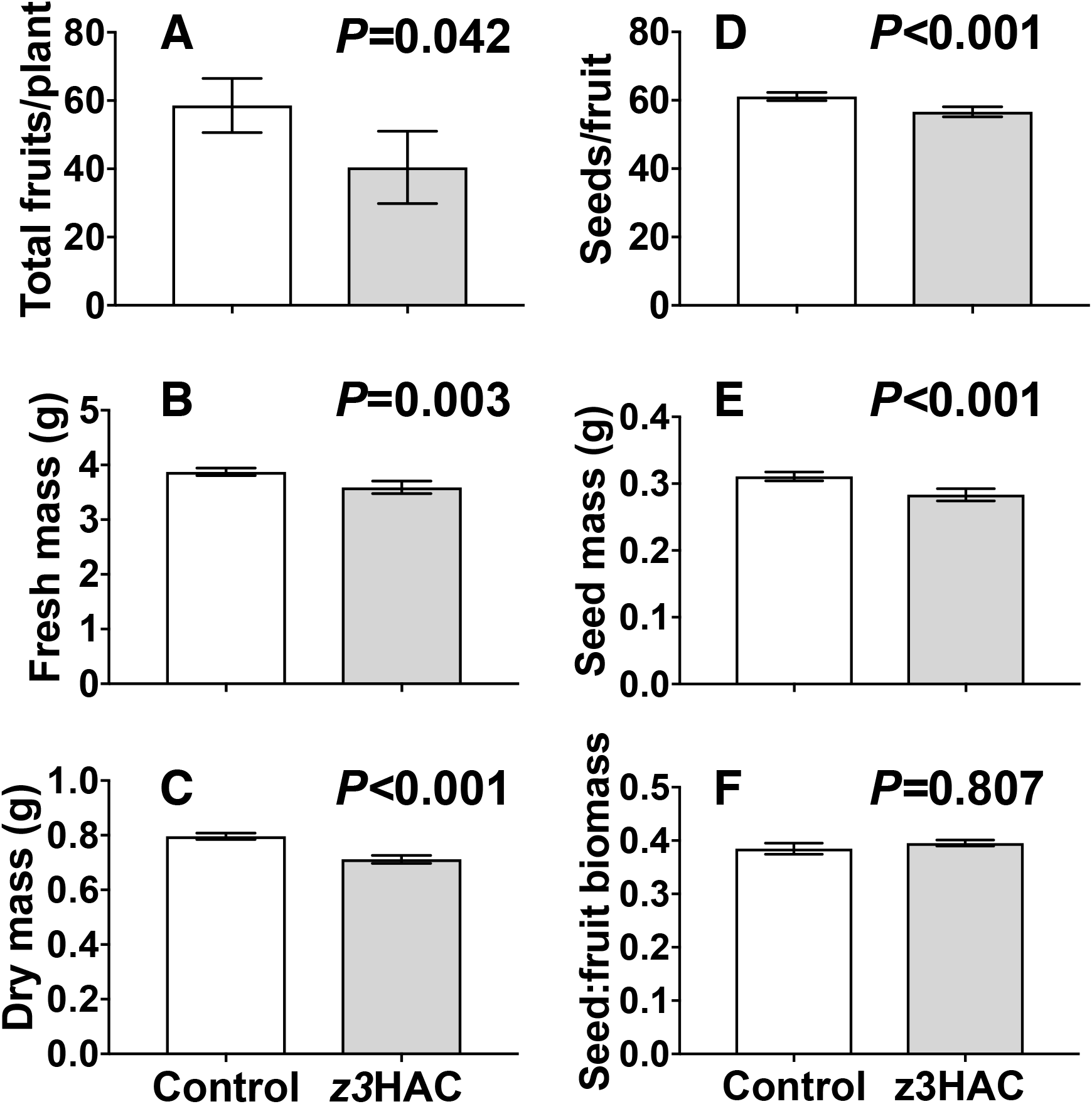
Fruit and seed production in *Capsicum annuum* (pepper) plants grown in a common garden experiment treated with a persistent application of the green leaf volatile *cis*-3-hexenyl acetate (*z*3HAC). The (A) total number of fruits were counted in the field, and (B) wet and (C) dry masses fruit masses were determined in the lab. (D) The total number of seeds per fruit and (E) the estimated mass per seed were determined from a subset of the fruits produced. (F) The ratio of seed mass to fruit mass was calculated to assess the efficiency of seed production. Bars represent means +/- S.E.M.. White boxes represent control plants; gray boxes represent plants treated with *z*3HAC. *P*-values represent Tukey’s HSD comparisons. See Supplemental Table 4 for complete statistics.

### z*3HAC exposure reduces herbivory on lima bean*

*z*3HAC exposure reduced natural herbivory in lima bean but not pepper plants. Chewing herbivory on pepper plants was low overall and statistically higher in *z*3HAC-treated plants only during the first sampling date (Fig.5A, F=_1,193_=5.627, *P=*0.019). In contrast, chewing damage to lima bean leaves increased as the field season progressed, with *z*3HAC-treated plants having overall 26% less chewing damage than did control plants (Fig.5B; F_1,539_=21.745, *P<*0.001). In addition to chewing herbivory, black bean aphids (*Aphis faba*) colonized 87% of the *z*3HAC-treated lima bean plants, compared with only 21% of control plants (Fig.5C; χ^2^ = 50.11, df=1, *P*<0.001). *A*.*faba* colonized early in the season and was only observed June 15-31 (Julian dates 166-181) because a heavy rainfall event reduced their population to undetectable levels. Piercing/sucking herbivores were rare for the remainder of the experiment.

**Fig. 5.**
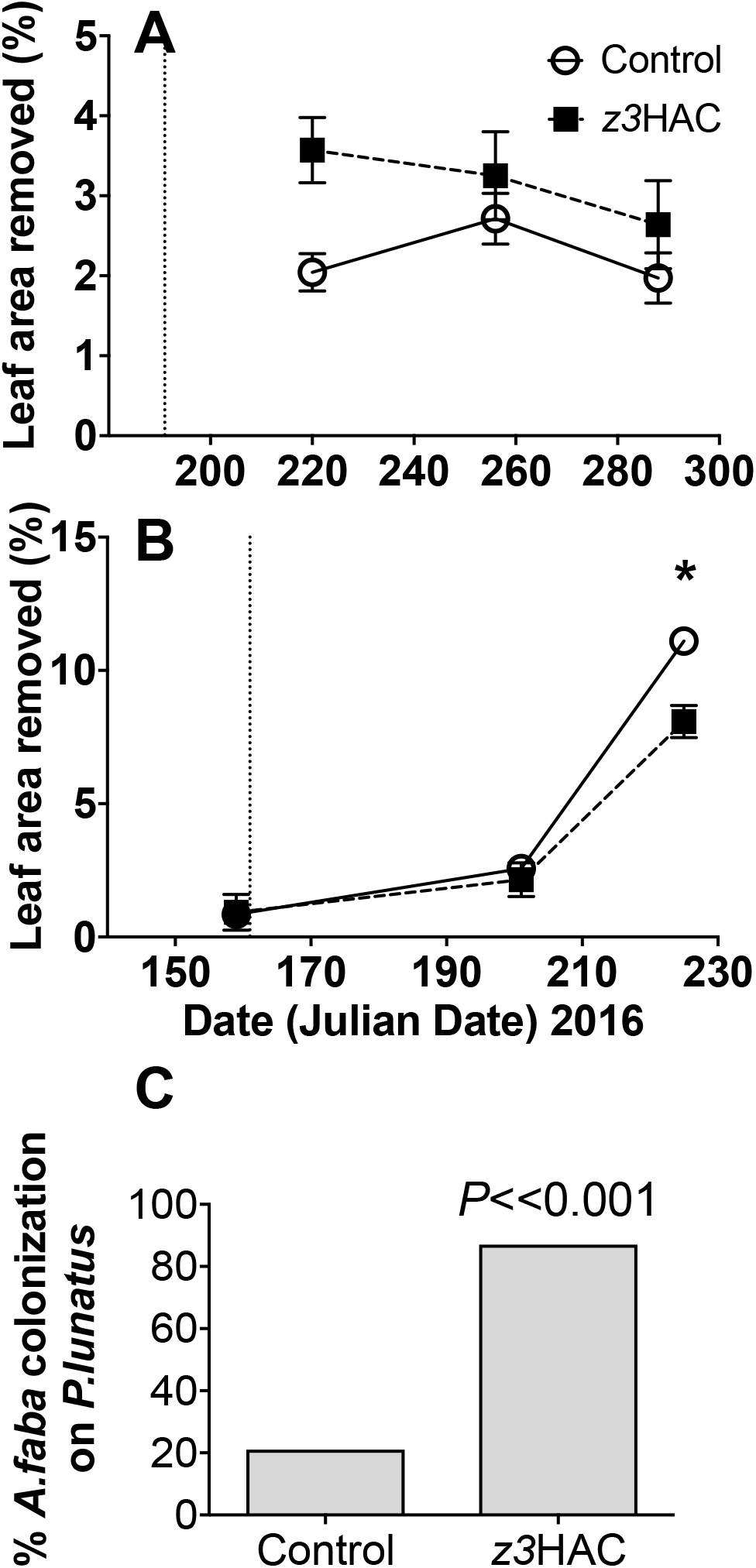
Herbivore damage on *Capsicum annuum* (pepper) and *Phaseolus lunatus* (lima bean) plants in a common garden field experiment. Chewing damage on (A) pepper and (B) lima bean plants was determined using a visual estimation technique (see Methods). Open circles represent control plants (receiving lanolin-filled vials); filled squares represent plants receiving a persistent application 10ng/hr *cis*-3-hexenyl acetate (*z*3HAC) dissolved in lanolin. Dropdown lines indicate the initial application of *z*3HAC treatment. Points represent averages +/- SE. Repeated measures ANOVAs (aov in R) were followed by one-way ANOVAs at each time-point. Asterisks (*) represent *P*<0.05 between treatment and control at each time point. See Supplemental Table 5 for complete statistics. (C) *Aphis faba* colonization on lima bean plants. Bars represent the percentage of plants in each group where *A*.*faba* were observed (χ^2^ = 50.11, df=1, *P*<0.001).

### Cyanogenic potential in lima bean is decreased by z3HAC exposure

Exposure to *z3*HAC reduced cyanogenic potential in lima bean plants. *z3*HAC exposure alone reduced cyanide concentration by 50% in sink leaves (Figure 6A; Z= -0.3257, *P=*0.003), but not significantly in source leaves (Fig. 6B; Z=-0.14387, *P=*0.146). Baseline cyanogenic potential was ∼2x greater in sink leaves compared to source leaves (Figure 6).

**Fig. 6.**
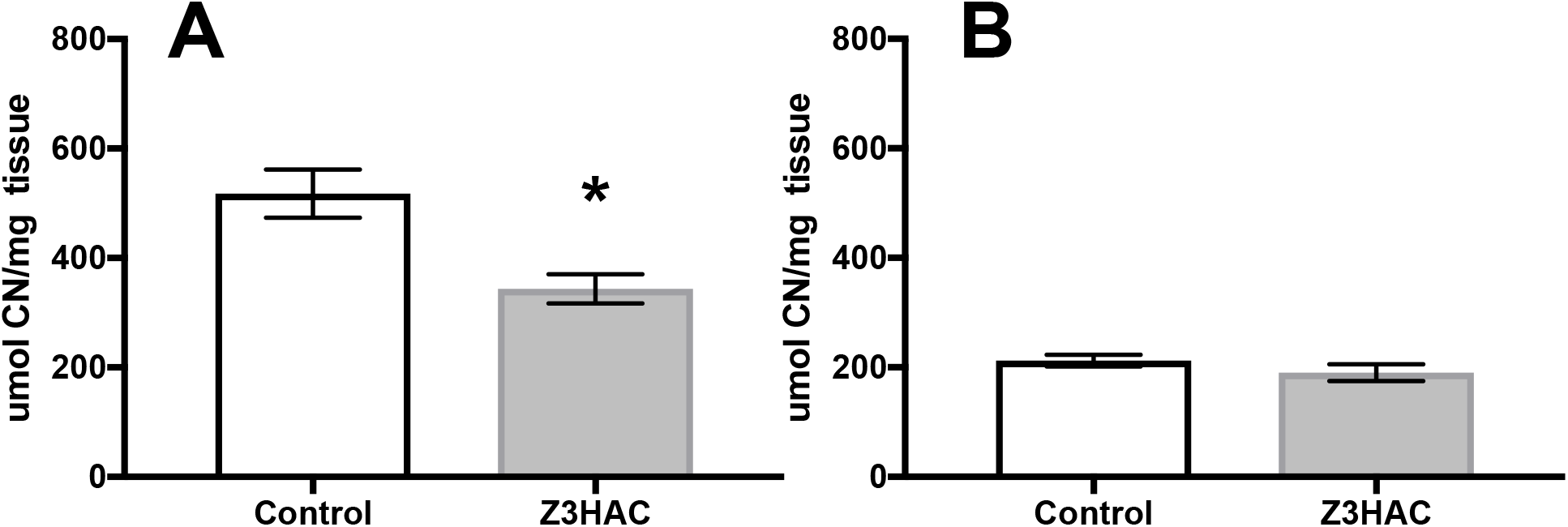
Cyanide induction in *Phaseolus lunatus* (Lima bean) as affected by *z3*HAC exposure. Cyanide concentration was determined by trapping volatile HCN in 1.0M NaOH followed by colorometric quantification. Sink leaves (A) and source leaves (B) were exposed long-term exposure of 10ng/hour *z3*HAC or left as a control. Points represent averages +/- SE and * indicate a p-value <0.05.

## Discussion

We show that a persistent, low-dose application of *z*3HAC differentially affects growth and reproduction of two plant species grown in the same field. Based on previous work on plant and sensory perception of volatiles [7,21], we hypothesized *z3*HAC application would decrease growth and reproductive fitness in both plant species. The rationale for this hypothesis was a central assumption of induced resistance theory that ecological costs modulate the deployment particular defensive phenotypes until necessary [30,31,56-60]. Volatile-mediated priming, even if regulated by a different mechanism from resistance [61], is an inducible phenomenon that theoretically should incur such fitness costs [36]. Yet, our results clearly indicate that pepper and lima bean had divergent fitness outcomes when subjected to a single GLV under identical field conditions. Whereas *z3*HAC-treated pepper plants had reduced growth (Fig. 1) and no effect on herbivore resistance (Fig. 5A) relative to controls, *z3*HAC-treated lima bean plants grew more and produced more flowers (Fig. 1,3), and suffered less chewing herbivory (Fig. 5B) compared to controls. This result—that some plants experience costs while others have minimal or even positive effects when exposed to the same HIPV—has important implications for how volatile cues may structure interspecific competition and ecological communities. HIPVs alone may be sufficient to result in differential fitness effects among species. Moreover, the positive effects of *z3*HAC on lima bean growth and flowering opens agronomic opportunities to exploit plant volatiles to enhance both growth and pest resistance of important crop plants.

What might affect the response of plants to volatile exposure? One possibility is that the signal integrity of HIPVs varies among plant species, which influences plant sensory perception and the outcome of defense priming. That is, *z*3HAC provides different information to different plants. Previous work on the role of HIPVs in plant anti-herbivore resistance focused on priming-mediated defense with consistent results in wheat [62,63], corn [12,45], lima bean [8,11,64], tomato [65], blueberry [15], sagebrush [43], Arabidopsis [66] and poplar [4]. In contrast, we specifically focused on indicators of plant fitness in lima bean and pepper in a common garden experiment. *z*3HAC treatment alone increased growth and flowering in lima bean, while reducing growth and reproductive output in pepper (Fig. 3 and 5). This season-long evidence illustrates that informational integrity on an HIPV varies between these two plant species. Such divergent fitness effects from exposure to a single ubiquitous herbivore-associated cue underscore the potential for functional similarity in the mechanisms by which plants modulate responses to herbivory and volatile indicators of herbivory.

Flower and fruit production is a key component of plant fitness potential. We show that *z*3HAC treatment alone differentially affected flower production in lima bean and pepper (Fig. 3). Insect herbivory can increase or decrease floral production depending on the system and environmental conditions [67-69]. Whereas increased flower production is a strategy assumed to ameliorate fitness losses in the presence of an environmental stress [56,70], decreased flower production may be related to costs of chemically mediated defense [71]. Previous work with lima bean has demonstrated stress-mediated compensation [72]. Our result that *z*3HAC alone was sufficient to trigger increased flowering is consistent with this observation, suggesting that *z*3HAC alone may stimulate a long-term stress response similar to herbivory. Therefore, even though the mechanisms underlying *z*3HAC-mediated effects on flower and fruit production are not yet known, they may be similar to those induced by herbivory [67].

Resource allocation between different tissues is pivotal for growth, reproduction, and defense, and can be influenced by environmental stress. For example, direct herbivory alters resource allocation between aboveground tissue and belowground tissue [49,73,74], as does application of the anti-herbivore phytohormone jasmonic acid (JA) [75,76]. Volatile cues can also affect biomass allocation. For example, barley exposed to volatiles from unwounded neighboring plants of different cultivars increases root and leaf biomass [77], while exposure to volatiles decreases aboveground biomass in other systems [78,79]. In our case, volatile treatment reduced overall aboveground and belowground biomass in pepper, but did not appear to alter overall biomass allocation patterns. Simply put, *z*3HAC-treated pepper plants were smaller overall, and therefore produced fewer seeds.

Differential investment between growth and defense is vital for maximizing limited resources and, ultimately, fitness. Inducible defenses in plants against herbivores and pathogens modulate such growth/defense tradeoffs in a number of plant species [31]. In some cases, exposure to VOCs alone is sufficient to affect such tradeoffs. For example, sagebrush exposed to VOCs from damaged conspecifics had decreased growth [80], while HIPV-exposed tobacco had increased herbivore resistance but decreased seed set [38]. In our experiment, *z*3HAC exposure decreased cyanogenic potential in sink leaves (Fig. 6), which were 20-30% higher than in source leaves. These results suggest that persistent exposure to *z*3HAC led to allocation shifts from cyanogenic potential (a putative defense) towards growth (Fig. 1 and 3), as well as supporting ontogeny-mediated cyanogenic potential [52,81].. That said, *z*3HAC-treated plants also experienced less natural damage primarily on mature source leaves than did controls (Fig. 5, personal observations), suggesting that the herbivory was deterred by defense mechanism other than cyanogenic potential in source leaves.

Volatile cues may impact ecological communities in both expected and pleiotropic ways. HIPVs are well-established mediators of multi-trophic antagonistic and mutualistic interactions [82-84], and manipulations of chemical signals and volatile blends have been used for biological control in a wide range of systems [85,86]. For example, HIPV-infused sticky traps in a grape (*Vitis vinifera*) orchard differentially attracted lacewings, hoverflies, and parasitoids [87]. Exogenous GLV manipulation using “dispensers” under field conditions altered the arthropod community composition in maize [47]. In our study, *A*.*faba* were clearly and unexpectedly attracted to *z*3HAC-exposed plants (Fig. 5). Under glasshouse conditions, *A*.*faba* were repelled by *z*3HAC alone [88], which suggests that the cue that mediated attraction was not our treatment alone. It is tempting to speculate that aphid attraction combined with reduced chewing herbivory in lima bean may be reflective of *z*3HAC effects on JA and salicylic acid (SA) signaling, which would be consistent with a JA-SA tradeoff [89,90]. Ultimately, however, the utility of GLVs (or other VOCs) in field applications will depend on understanding community-level effects of volatile exposure.

As a caveat, volatile identity, concentration, and duration may affect the reliability of a cue and therefore the costs associated with eavesdropping. Plants experiencing insect herbivory frequently generate species-specific blends of volatile compounds [91,92], which can influence fitness in neighboring plants [80,93,94]. Plant-derived compounds associated with herbivory include GLVs [4,12,78], shikimate derivatives [13], and terpenes [95]. However, individual compounds within a blend can affect plant defense and priming as much as the blend itself. We used *z*3HAC in this study because it is released herbivore-damaged leaves [21], which ostensibly allows *z*3HAC to confer reliable ecological information. That said, *z*3HAC can also be released from wounded leaves without herbivores [91] [23]. However, plants detect and respond specifically to *z*3HAC [4], and the costs associated with that response in the context of pest resistance were the focus of this investigation. Additionally, concentration of a cue may influence plant resistance [66,78]. For example, a repeated, low-dose exposure to a GLV blend enhances plant resistance compared to a single application [46] while *z3*HAC emissions can be as high as 66 ng/cm^3^ after herbivory [20]. For these reasons, we chose to use a low-dose exposure to *z*3HAC (25% of the concentration that primed poplar [4] and maize [21]), and still observed divergent fitness effects between the two plant species (Figs.1, 3, and 5).

In summary, our key finding is that persistent application of a low dose of a single volatile compound *z*3HAC, a common HIPV and GLV, in field conditions leads to divergent growth and reproductive fitness effects between two plant species. This result underscores the variable nature of volatile-mediated eavesdropping, and that plants may have evolved species-specific mechanisms for responding to volatile cues. Given natural variation among species, future work must assess costs with other species as well as within accessions and landraces of our model plants over multiple years. Ultimately, the adaptive significance of eavesdropping for enhancing plant immunity will depend on plant life history, physiology, and other ecological factors to determine whether a plant will benefit from eavesdropping VOCs or not, and therefore what impact volatile-mediated eavesdropping might have on plants and their insect pests.

## Supplemental Material

Supplemental Tables describing the outputs of statistical analyses are listed as tabs in a single spreadsheet (“Supp_Tables”), with tab names corresponding to each table. Supplemental Figure 1 (“Supp_Fig1”). Aerial view of the field site within the Blackacre Community Garden. The research area is outlined and the original picture was acquired from Google Earth. Supplemental Figure 2 (“Supp_Fig2”). Volatile administration setup. Each 2mL glass vial was inverted and supported by wire stands to prevent rain water accumulation.

Additionally, each vial was wrapped in aluminum foil to prevent photodegradation [47]. Supplemental Figure 3 (“Supp_Fig3”). Calculated individual seed mass for *C*.*annuum* (A) and total pod production for *P*.*lunatus* (B). Box plots represent the raw data ranging from the upper to the lower quartiles and the median. White boxes represent control plants; gray boxes represent plants treated with *z*3HAC. Error bars represent the 5% and 95% of the data, and individual dots are observations that fell outside of those parameters. *P*-values represent Tukey’s HSD comparisons.

## Acknowledgments

We are grateful to Andrea Clavijo-McCormick for the invitation to submit our work to this special issue. We thank Allie Peot and Abhinav Maurya for field assistance, and to A.Peot for assistance processing the plant materials in the laboratory. Comments from Heidi Appel, Amy Austin, and anonymous reviewers greatly improved the manuscript. We are grateful to A. Dale Josey and Susan Ballerstedt for permission and logistical support to work in the community gardens at the Blackacre Conservancy.

