## Supplementary figures and images for "Dispensing a synthetic green leaf volatile to two plant species in a common garden differentially alters physiological responses and herbivory"

### Supplemental Figure 1

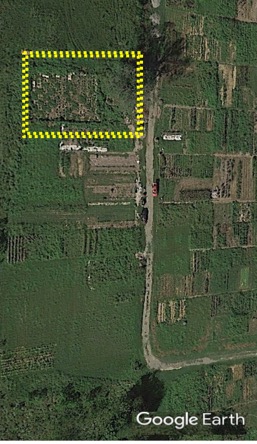

### Supplemental Figure 2

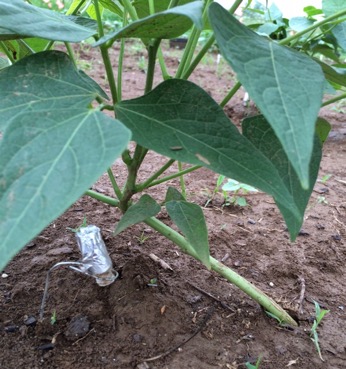

### Supplemental Figure 3

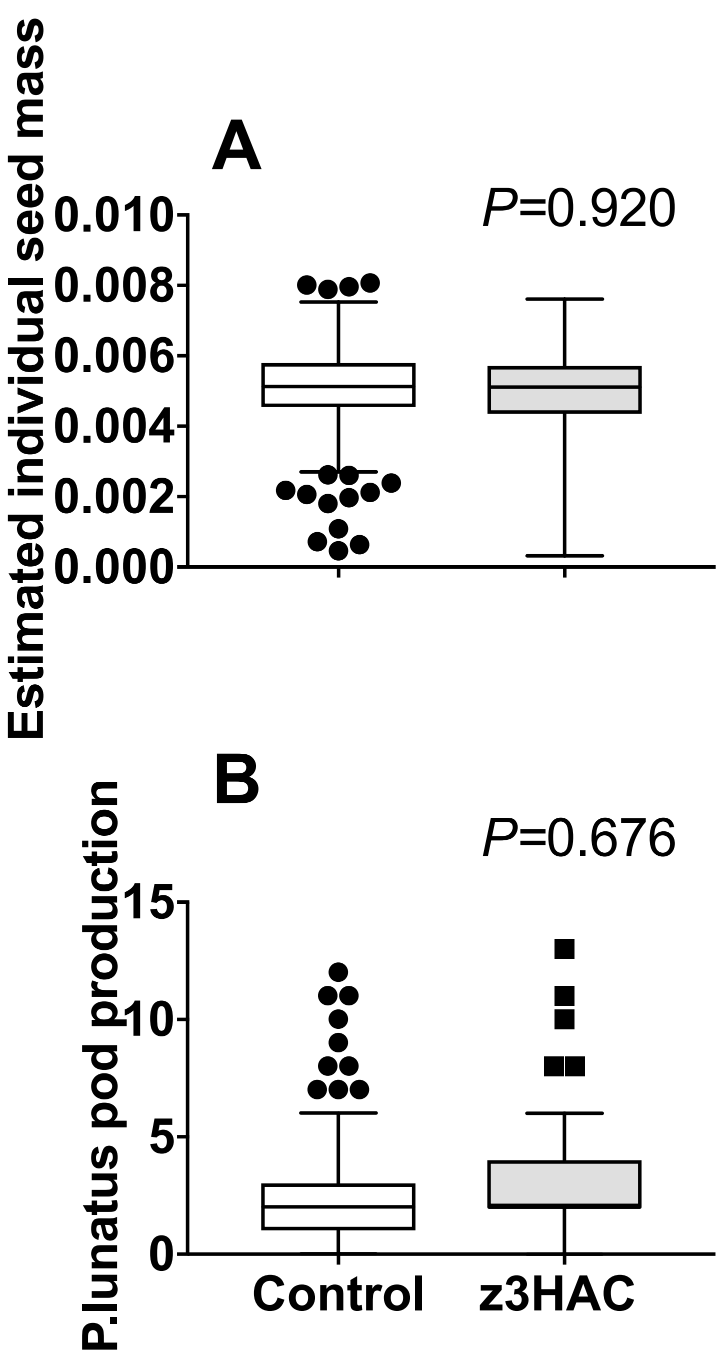
